# Perinatal Shifts in Fecal-Derived Metabolites and Associations with Postpartum Depression

**DOI:** 10.1101/2025.09.25.678627

**Authors:** Kayla D. Longoria, Michelle L. Wright, Kimberly A. Lewis, Sarina R. Garcia, Oscar Y. Franco-Rocha, Sara Dube, Tien C. Nguyen, Elena Flowers, Elizabeth M. Widen

## Abstract

**Introduction:** Research on maternal depression is largely limited to static, blood-derived biomarkers in the postpartum period and mechanistic targets derived from populations outside the physiological contexts of pregnancy and postpartum, resulting in critical gaps in understanding context-specific mechanisms underlying this debilitating condition.

**Objectives:** To examine temporal shifts in the maternal gut metabolome and associations between pregnancy-specific shifts and postpartum depression (PPD).

**Methods:** We applied untargeted metabolomics (UPLC-MS/MS) to fecal samples collected from participants (N= 25) enrolled in the Maternal and Infant NutriTion (MINT) study. Random forest analysis was used to identify key pathways and metabolites contributing to temporal shifts. Fold change analysis and paired t-tests were used to quantify the magnitude and significance of metabolite changes.

Associations between pregnancy-specific changes and PPD (Edinburgh Postnatal Depression Scale at 6 weeks) were identified using Pearson’s correlation.

**Results:** Lipid, amino acid, and xenobiotic metabolism emerged as core pathways driving temporal changes in the maternal gut metabolome. The most pronounced shifts occurred from 35 weeks gestation to postpartum, with 55 metabolites significantly altered compared to 24 from 24-to 35 weeks gestation and 26 from 24 weeks gestation to postpartum. Of the 29 metabolites associated with PPD; 68.9% were metabolic intermediates, primarily involved in lipid and amino acid metabolism (58.6%).

**Conclusions:** This study provides some of the first evidence of temporal shifts in the maternal gut metabolome and associations with PPD, highlighting the importance of lipid and amino acid metabolism and laying the groundwork for future multi-omics research.

## BACKGROUND

Postpartum depression (PPD) is a prevalent and debilitating mental health condition that affects up to 37% of birthing individuals (Dagher et al., 2021; Van Niel & Payne, 2020). The consequences of PPD extend beyond the birthing individual, impacting the short-and long-term health of children and contributing to an estimated economic burden of $7.5B per each year of births (Luca et al., 2019, 2020). Metabolic activity during pregnancy is understood to function from a baseline distinct from that of non-pregnant states (Kepley, 2023). While these shifts are essential to maintain maternal-fetal homeostasis, they impose prolonged physiological demands that can strain the body’s adaptative capacity, potentially compromising the health of both mother and child. Dysregulated metabolic activity during pregnancy is evidenced to contribute to various pathophysiological conditions (i.e., gestational diabetes, preeclampsia), with residual consequences extending into the postpartum period (Dudzik et al., 2014; Hauspurg & Jeyabalan, 2022; Ott et al., 2020; Ross et al., 2019; Spanou et al., 2022; Toloza et al., 2022). Despite this and evidence indicating up to 80% of PPD cases report symptom onset begins in pregnancy (Cox et al., 2016; Wilcox et al., 2021), studies examining the relationship between pregnancy-related metabolic activity and PPD remain rare (Longoria et al., 2024).

Recent Zuranolone trials, leading to the first FDA approved oral PPD treatment, indicate that PPD arises from dysregulated communication within a biological network involving the endocrine system, neurosteriods, and neurotransmitter pathways (Deligiannidis et al., 2021, 2023). The gut microbiome, comprised of a diverse community of microorganisms (microbes) that reside in the gastrointestinal tract, work symbiotically with their host (human) to support essential physiological processes, such as metabolism, immune regulation, and neuroactive signaling (de Vos et al., 2022; He et al., 2021; Shoubridge et al., 2022). During pregnancy and postpartum (perinatal), the gut microbiome undergoes significant shifts in response to the metabolic demands induced by these two periods, suggesting it likely serves as a key regulator hub that facilitates communication between biological systems (gut-brain-axis) for the purposes of maintaining maternal-child homeostasis (Sinha et al., 2023; Weerasuriya et al., 2023). However, the functional implications of these shifts remain unclear in perinatal populations, as studies have largely been limited to gut microbial composition rather than its biochemical output (metabolites)—the functional intermediates (bioactive compounds) and byproducts (waste) that provide insight into microbiome-host interactions.

Though a few studies have examined interactions between blood-derived metabolites (systemic) and the gut microbiome (local) in the context of PPD (Cui et al., 2024; Zhang et al., 2023), blood-derived metabolites primarily reflect heavily processed metabolites that have undergone extensive modification in the liver and other organs, making it difficult to identify early, localized microbial contributions in metabolic regulation and potential dysregulation (Doan et al., 2024; van de Graaf et al., 2024). Fecal-derived metabolites, however, offer a proximal view of microbial activity before systemic absorption (Deng et al., 2023; Donatti et al., 2020; Xu et al., 2019), positioning them as particularly valuable for understanding microbiome-host interactions that may foreshadow metabolic dysregulation and adverse health outcomes postpartum (e.g., PPD). This distinction is particularly crucial in the context of perinatal depression, as maternal metabolic activity influences offspring health and development (Adank et al., 2020; Parrettini et al., 2020) and oral psychopharmacological interventions (e.g., selective serotonin reuptake inhibitors (SSRIs), Zuranolone) are first-line interventions when psychotherapy is inaccessible, ineffective, or insufficient (Cox et al., 2016; Kaufman et al., 2022).

Addressing critical gaps in understanding how adaptations in metabolic activity during pregnancy influence PPD risk will enable the discovery of novel risk biomarkers with high translational potential, as pregnancy is a period of increased healthcare engagement. By applying untargeted metabolomics to maternal fecal samples collected during pregnancy and postpartum, this study will provide some of the first evidence of longitudinal changes in the maternal gut metabolome and associations between pregnancy-related changes and PPD symptom severity. These findings will inform future multi-omics studies integrating localized (gut) and systemic biomarkers to advance understanding of the biological networks underlying PPD risk, as well as other perinatal health conditions.

## METHODS

### Design

This study utilized data from the Maternal and Infant Nutrition (MINT) study (NCT04132310). The MINT study is an ongoing prospective observational cohort study following mother-child dyads to examine relationships between pregnancy weight trajectories and maternal-child health outcomes postpartum. Beginning in August 2019, adult (≥ 18 years) pregnant women <16 weeks gestation were recruited from obstetrics and gynecology (OB/GYN) clinics in two commonly utilized hospital systems located in Central Texas. Participants were excluded from participation if they had a body mass index (BMI) > 35kg/m2, diabetes or any other medical history that may influence weight outcomes, or any magnetic resonance imaging (MRI) contraindications.

Leveraging maternal fecal samples collected at 24-and 35 weeks gestation and 6 weeks postpartum, the current study collected *de novo* untargeted metabolomics data from a subset of participants (N = 25) enrolled between August 2019 – June 2022. Sociodemographic characteristics were collected at baseline (≤16 weeks gestation) and PPD symptom severity was measured at 6 weeks postpartum. Participants included in the present analysis were those who had complete datasets for all time points by June 2022, which is when the fecal samples underwent processing for subsequent analyses. This study was approved and monitored by the Institutional Review Board at The University of Texas at Austin (2018-05-0127).

### Measures

Sociodemographic data were collected via self-report questionnaires. PPD symptom severity was measured by the Edinburgh Postnatal Depression Scale (EPDS) at 6 weeks postpartum using total scores as a continuous variable. The EPDS is a 10-item self-report questionnaire widely used in clinical practice and research to screen for PPD symptoms (Cox et al., 1987; Levis et al., 2020; Moyer et al., 2023). It uses a 4-point Likert scale (0-3) to measure feelings of depression over the last 7 days, with varying sensitivity (59-100%) and specificity (49-100%). The total score ranges from 0-30, with higher scores indicating greater symptom severity.

Fecal samples were collected in sterile cryovials at 24-and 35 weeks gestation and 6 weeks postpartum following standardized protocols to maintain sample integrity and compliance with all ethical and regulatory standards. Samples were stored at-80°C in a restricted access freezer at The University of Texas at Austin until further analyses. In June of 2022, samples were removed from the freezer, and 100mg of solid material was aliquot and shipped on dry ice to Metabolon, Inc. (Morrisville, NC, USA) for untargeted metabolomics analysis. Methods used by Metabolon for metabolite data acquisition and analyses are described below (Metabolon Inc., 2023).

### Metabolite data acquisition

Upon receipt, samples were inventoried and stored at-80°C. Samples were accessioned into the Metabolon Laboratory Information Management System (LIMS) and assigned a unique identifier linked only to the original source identifiers. This identifier was used to track all sample handling, tasks, and results, with new identifiers automatically assigned for each processing step.

#### Sample preparation

Samples were prepared using the automated MicroLab STAR® system (Hamilton Company). Recovery standards were added before extraction for quality control. To recover chemically diverse metabolites, proteins were precipitated with methanol under vigorous shaking for 2 mins (Glen Mills GenoGrinder 2000) followed by centrifugation. The resulting extract was divided into five fractions: two for analysis by two separate reverse phase/ultrahigh performance liquid chromatography-tandem mass spectroscopy (RP/UPLC-MS/MS) methods with positive ion mode electrospray ionization (ESI), one for analysis by RP/UPLC-MS/MS with negative ion mode ESI, one for analysis by hydrophilic interaction liquid chromatography (HILIC)/UPLC-MS/MS with negative ion mode ESI, and one sample was reserved for backup. Organic solvent was removed using TurboVap® (Zymark), and extracts were stored overnight under nitrogen prior to preparation for analysis.

#### Quality assurance/Quality control (QA/QC)

Multiple controls were analyzed alongside study samples. A pooled matrix sample, created by combining small volumes from each study sample, served as a technical replicate. Extracted water samples were used as blanks. A proprietary QC standard cocktail designed to avoid interference with endogenous compounds was spiked into all samples to monitor instrument performance and aid chromatographic alignment. Instrument variability was determined by the median standard of the spiked standards, while overall process variability was determined in all pooled matrix samples. Study samples were randomized, with QC samples evenly distributed throughout the platform run.

#### Ultrahigh performance liquid chromatography-tandem mass spectroscopy (UPLC-MS/MS)

Samples were analyzed using a Waters ACQUITY UPLC system and a Thermo Scientific Q-Exactive high-resolution/mass spectrometer interfaced with a heated electrospray ionization (HESI-II) source and Orbitrap mass analyzer operated at 35,000 mass resolution. Extracts were dried and reconstituted in solvents compatible to each of the four methods. The four analytic methods were: (1) hydrophilic compounds were analyzed in an acidic positive ion condition using C18 column (Waters UPLC BEH C18-2.1×100 mm, 1.7 µm) with water and methanol, containing 0.05% perfluoropentanoic acid and 0.1% formic acid; (2) hydrophobic compounds were analyzed in an acidic positive ion condition using C18 column with methanol, acetonitrile, water, 0.05% perfluoropentanoic acid and 0.01% formic acid and were operated at an overall higher organic content; (3) basic compounds were analyzed in negative ion mode using methanol and water, however with 6.5mM Ammonium Bicarbonate at pH 8; (4) polar compounds were analyzed in negative ion mode using HILIC column (Waters UPLC BEH Amide 2.1×150 mm, 1.7 µm) with water and acetonitrile with 10mM Ammonium Formate (pH 10.8). Mass spectrometry alternated between MS and data-dependent MS^n^ scans with dynamic exclusion, covering 70-1000 m/z. Raw data files were archived and extracted.

#### Metabolite identification and quantification

Raw data were extracted, peak-identified, and QC-processed using Metabolon’s proprietary hardware and software. Compounds were identified by comparison to library entries of purified standards (> 3300) or recurrent unknown entities. Biochemical identifications were based on three criteria: (1) retention index within a narrow window of the proposed identification; (2) accurate mass match to the library +/-10 ppm; and (3) the MS/MS forward and reverse scores between the experimental data and authentic standards. Peaks were quantified using area-under-the-curve.

### Statistical analysis

Descriptive statistics were used to summarize participant characteristics, with means and standard deviations (SDs) for continuous variables and frequencies and percentages for categorical variables. Metabolite data were log transformed prior to analyses. Since psychiatric history is a well-established risk factor of PPD (Liu et al., 2022; Yang et al., 2022), Principal Component Analysis (PCA) was applied to evaluate differences in metabolic profiles between women with a self-reported mood disorder history (i.e., depression, anxiety) at some point in their lifetime compared to those without. Random Forest Analysis was used to identify key metabolites and metabolic pathways that most strongly contributed to metabolic shifts across perinatal timepoints. Fold change (FC) analysis and paired t-tests were used to quantify the magnitude and significance of metabolite changes across perinatal timepoints. Pearson’s correlation was applied to evaluate associations between metabolite changes in pregnancy and PPD symptom severity. Analyses were performed in ArrayStudio/Jupyter Notebook and R (version 4.2.2), with α set at < 0.05 for significance. Adjustments for multiple comparisons were applied using False Discovery Rate (FDR), with a threshold of q < 0.05 (Benjamini & Hochberg, 1995). Aside from sertraline findings, only metabolites detected in ≥ 80% of samples across timepoints are discussed.

## RESULTS

### Participant characteristics

Participants were primarily Non-Hispanic (76%) White (80%) women with a mean (SD) age of 33.5 (4.3) years (**Table 1)**.

**Table 1.**
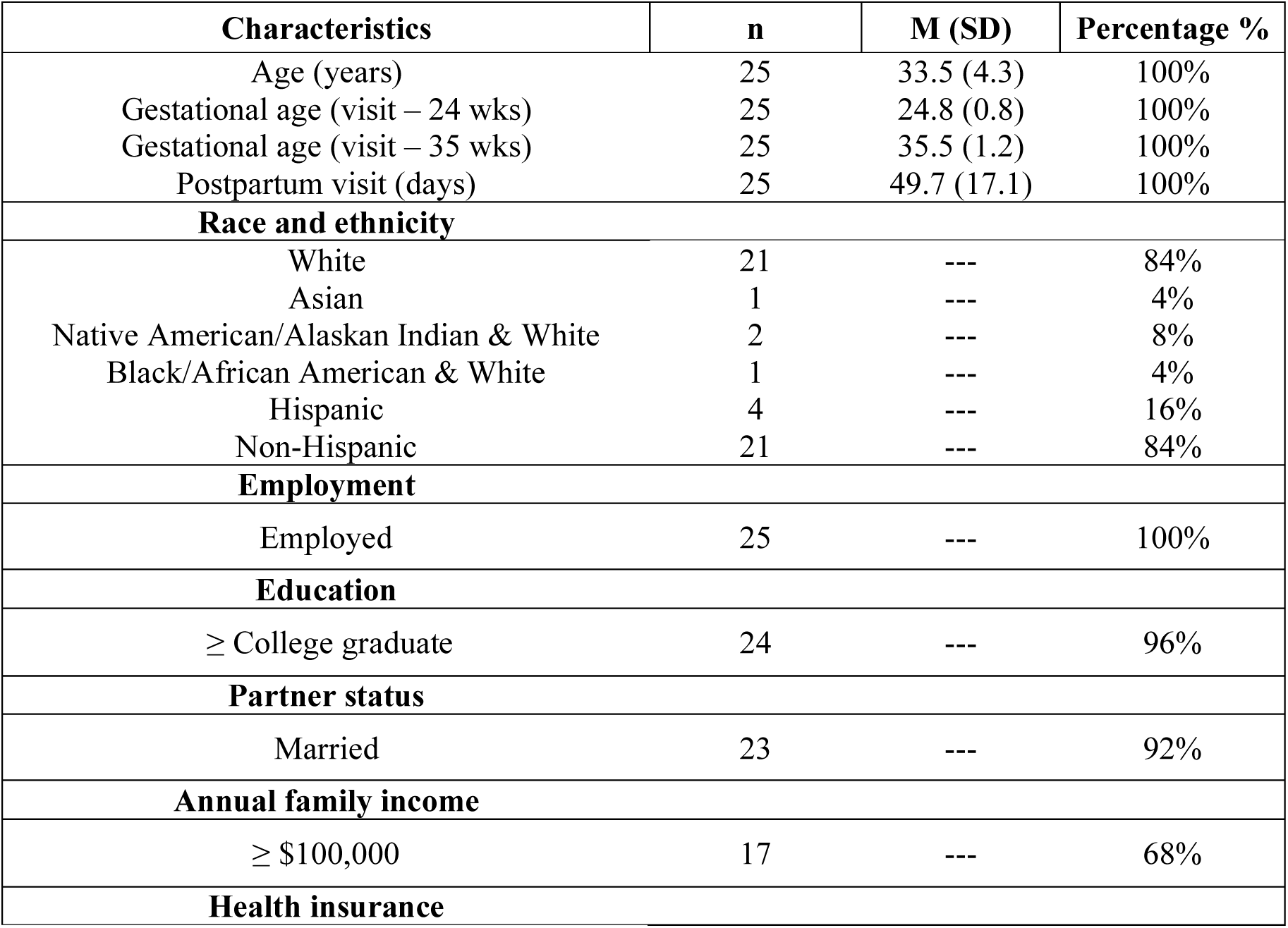
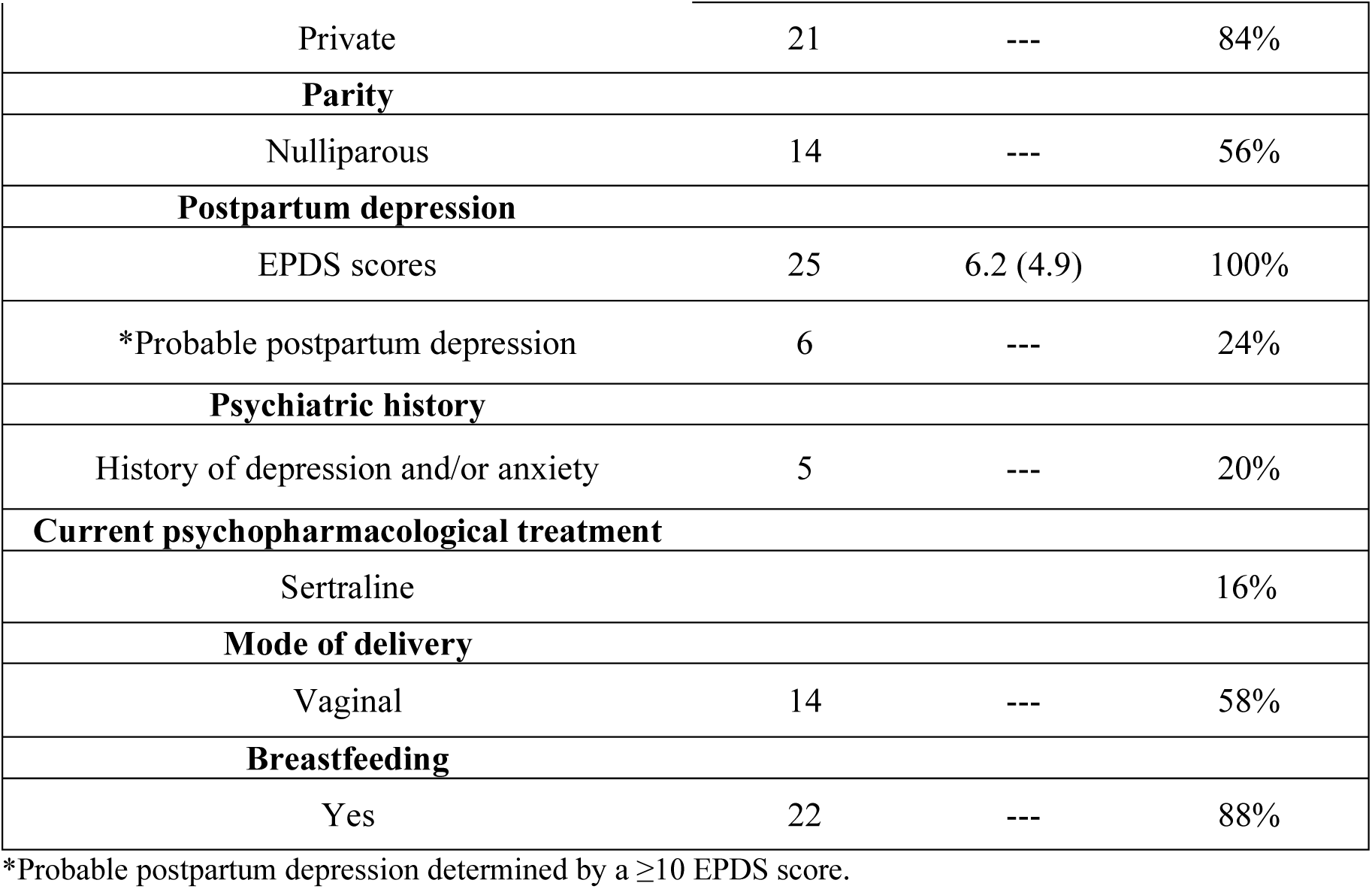
Participant characteristics.

| Characteristics | n | M (SD) | Percentage % |
| --- | --- | --- | --- |
| Age (years) | 25 | 33.5 (4.3) | 100% |
| Gestational age (visit – 24 wks) | 25 | 24.8 (0.8) | 100% |
| Gestational age (visit – 35 wks) | 25 | 35.5 (1.2) | 100% |
| Postpartum visit (days) | 25 | 49.7 (17.1) | 100% |
| **Race and ethnicity** |  |  |  |
| White | 21 | --- | 84% |
| Asian | 1 | --- | 4% |
| Native American/Alaskan Indian & White | 2 | --- | 8% |
| Black/African American & White | 1 | --- | 4% |
| Hispanic | 4 | --- | 16% |
| Non-Hispanic | 21 | --- | 84% |
| **Employment** |  |  |  |
| Employed | 25 | --- | 100% |
| **Education** |  |  |  |
| ≥ College graduate | 24 | --- | 96% |
| **Partner status** |  |  |  |
| Married | 23 | --- | 92% |
| **Annual family income** |  |  |  |
| ≥ \$100,000 | 17 | --- | 68% |
| **Health insurance** |  |  |  |
| Private | 21 | --- | 84% |
| **Parity** |  |  |  |
| Nulliparous | 14 | --- | 56% |
| **Postpartum depression** |  |  |  |
| EPDS scores | 25 | 6.2 (4.9) | 100% |
| *Probable postpartum depression | 6 | --- | 24% |
| **Psychiatric history** |  |  |  |
| History of depression and/or anxiety | 5 | --- | 20% |
| **Current psychopharmacological treatment** |  |  |  |
| Sertraline |  |  | 16% |
| **Mode of delivery** |  |  |  |
| Vaginal | 14 | --- | 58% |
| **Breastfeeding** |  |  |  |
| Yes | 22 | --- | 88% |

\*Probable postpartum depression determined by a $\geq 10$ EPDS score.

An EPDS score of ≥ 10 classified 24% of the sample as likely experiencing clinical PPD, falling within the incidence range commonly observed in the broader perinatal population (Dagher et al., 2021; Van Niel & Payne, 2020). Five participants (20%) reported having a mood disorder at some point in their lifetime. At baseline, four (16%) reported using a psychopharmacological intervention (i.e., sertraline) to treat a current mood disorder.

### Metabolic profiles differ by mood disorder history

The PCA revealed distinct clustering of metabolic profiles based on self-reported mood disorder history **(Figure 1).**

**Fig 1.**
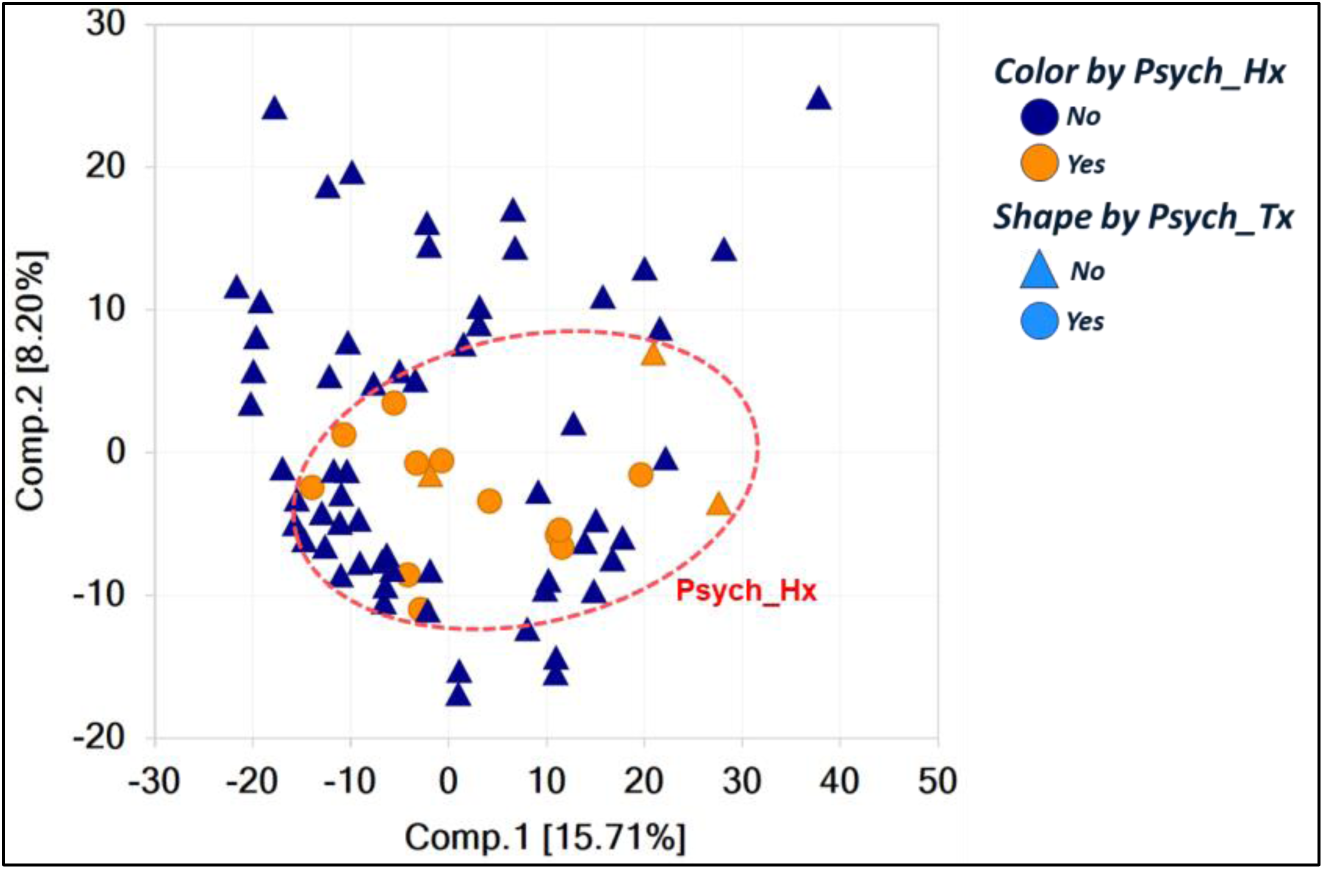
Principal Component Analysis of metabolite profiles by mood disorder history and current treatment.

Participants reporting a mood disorder history exhibited reduced metabolic variability, indicating greater similarity in metabolite profiles compared to those without. Those receiving psychopharmacological treatment at baseline did not exhibit any distinct clustering, indicating treatment status did not significantly contribute to variance in metabolic profiles. However, the absence of distinct clustering based on treatment status may be attributable to the small number of participants (n = 4) receiving treatment and/or variability in the duration or efficacy of the treatment used.

### Contributors to Perinatal Shifts in Fecal-derived Metabolites

Fecal-derived metabolites were modestly successful in distinguishing between pregnancy and postpartum timepoints, with an overall predicative accuracy of 60%. A random forest confusion matrix indicated postpartum samples were classified with the highest accuracy (96%, class error = 0.04), whereas pregnancy timepoints (24-and 35 weeks gestation) exhibited greater misclassification, with accuracies of 49% (class error = 0.51) and 33% (class error = 0.67), respectively **(Fig 2A)**. The biochemical importance plot identified three key metabolic pathways and 30 fecal-derived metabolites that most strongly contributed to metabolic shifts across perinatal timepoints, many of which were hormones. The most prominent pathways contributing to metabolic shifts between timepoints were lipid metabolism (19 metabolites), xenobiotic metabolism (7 metabolites), and amino acid metabolism (3 metabolites) **(Fig 2B)**. Individual metabolites, biological role(s), and empirically supported genes associated with the identified metabolites are detailed in **Table 2**.

**Fig 2.**
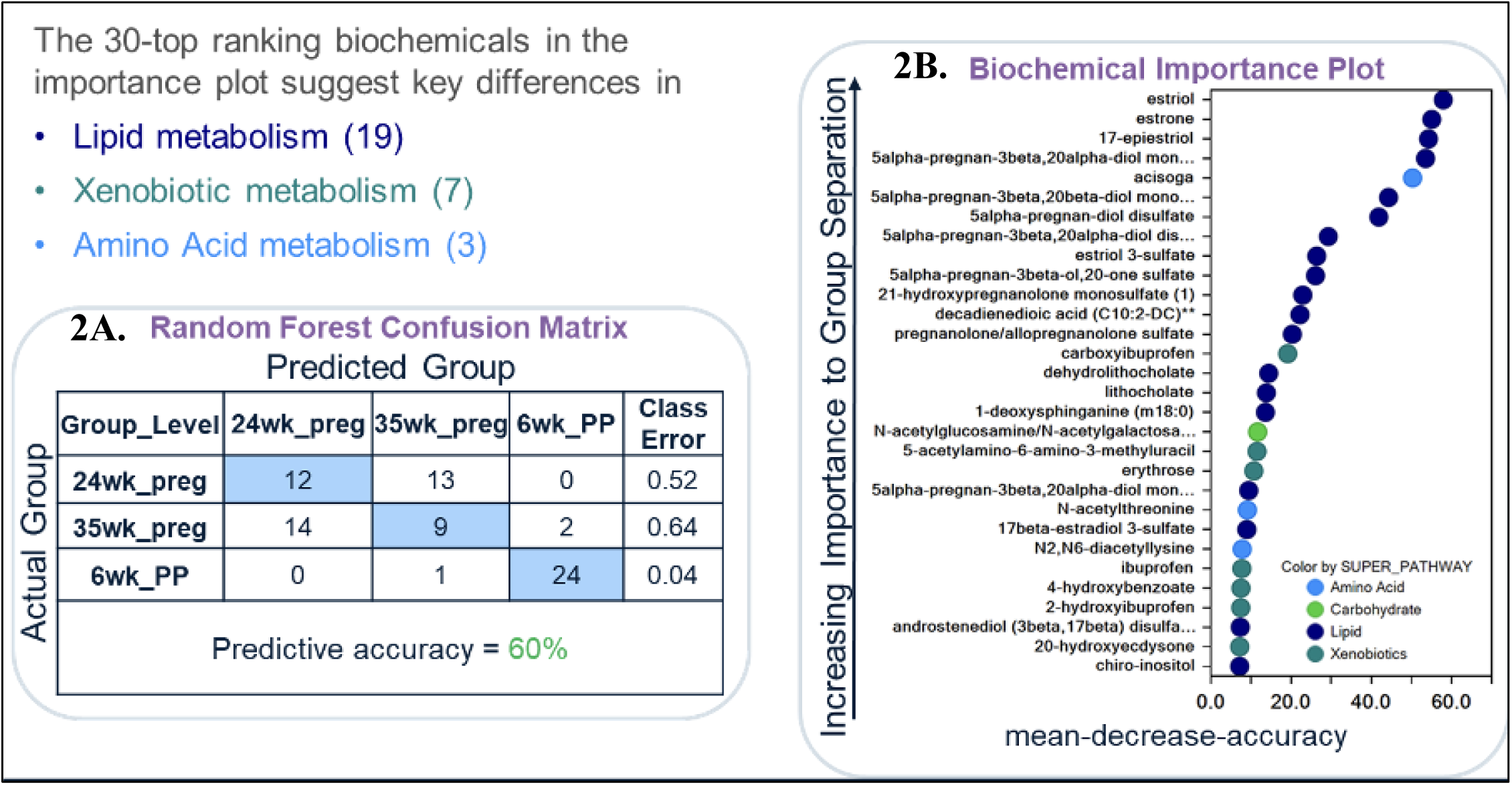
Key Contributors to Perinatal Shifts in Fecal-derived Metabolites. The Random Forest Confusion Matrix shows the predictive accuracy of fecal-derived metabolites in distinguishing between perinatal timepoints (Figure 2A). The Biochemical Importance Plot illustrates core pathways and metabolites contributing to metabolic shifts between perinatal timepoints (Figure 2B).

**Table 2.** Top 30 gut metabolites contributing to differences in perinatal timepoints Metabolites are listed in descending order as determined by the Biochemical Importance Plot (Figure 2B). Empirically derived information was acquired from The Human Metabolome Database (HMDB) (https://www.hmdb.ca/), as well as existing literature, indicated by PMID when HMDB did not provide any information for a specific metabolite.

|  |  |  | **Empirically derived** |  |  |
| --- | --- | --- | --- | --- | --- |
|  | **Gut Metabolite**<br>**(HMDB ID)** | **Pathway,**<br>**Subpathway** | **Detection**<br>**locations** | **Physiological**<br>**effect(s)** | **Gene family [n] - role** |
| 1 | Estriol<br>(HMDB0000153) | Lipid,<br>Estrogenic<br>Steroids | • Amniotic fluid<br>• Blood<br>• Urine | • Pregnancy<br>• Type 2 Diabetes (T2D) | **1.UGT** [18] - glucuronidation<br>**2.HSD17B** [4] - steroid metabolism<br>**3.ESR** [3] – estrogen signaling |
| 2 | Estrone<br>(HMDB0000145) | Lipid,<br>Estrogenic<br>Steroids | • Blood<br>• Liver<br>• Kidney<br>• Placenta<br>• Urine<br>• Hepatic tissue | • Major depressive disorder<br>• Vomiting<br>• Nausea<br>• Irritability<br>• Edema<br>• Bleeding | **1.UGT** [20] – glucuronidation<br>**2.HSD17B** [6] – steroid metabolism<br>**3.CYP (Cytochrome P450)** [21] – hormone and drug metabolism<br>**4.SULT** [1] – sulfonation<br>**5.STS** [1] – steroid sulfatase<br>**6. ESR** [3] – estrogen signaling<br>**7.Other hormone and transport-related genes [4]**<br>SHBG – sex hormone-binding globulin<br>ALB – Albumin carrier protein<br>SLC22A7 – organic anion transporter |
|  |  |  |  |  | 8. **ARSD/ARSE** [2] - Arylsulfatases |
| 3 | 17-epiestriol<br>(HMDB0000356) | Lipid,<br>Estrogenic<br>Steroids | • Urine<br>• Hepatic Tissue | None reported | 1. **ESR** – estrogen signaling (PMID: 12547825) |
| 4 | 5alpha-pregnan-3beta,20alpha-diol monosulfate (1)<br>(No dedicated entry, 1 measurement identified in GWAS catalog) | Lipid, Progestin Steroids | None reported | None reported | No currently published evidence directly associating metabolite with specific human genes. |
| 5 | Acisoga<br>(HMDB0061384) | Amino Acid, Polyamine Metabolism | • Urine<br>• Feces | • Type 2 Diabetes (PMID: 32751974) | 1. **ABDH3** – lysine metabolism (PMID: 32751974) |
| 6 | 5alpha-pregnan-3beta,20beta-diol monosulfate (HMDB0240580)<br>“very few articles have been published” | Lipid, Progestin Steroids | • Placenta | • Spontaneous preterm birth (PMID: 32033212) | No currently published evidence directly associating metabolite with specific human genes. |
| 7 | 5alpha-pregnan-diol disulfate (HMDB0240581, HMDB0240582) | Lipid, Progestin Steroids | • Placenta | • Spontaneous preterm birth (PMID: 32033212) | No currently published evidence directly associating metabolite with specific human genes. |
| 8 | 5alpha-pregnan-3beta,20alpha-diol disulfate (HMDB0094650) | Lipid, Progestin Steroids | • Blood<br>• Feces<br>• Urine<br>• Placenta | • Colorectal cancer (PMID: 27275383)<br>• Spontaneous preterm birth (PMID: 32033212) | No currently published evidence directly associating metabolite with specific human genes. |
| 9 | Estriol 3-sulfate<br>(No detected entry, PubChemID 66415) | Lipid, Estrogenic Steroids | • Plasma, Urine (PubMed list) | • Pregnancy | No currently published evidence directly associating metabolite with specific human genes. |
|  |  |  | Breast cyst fluid (PMID: 1387652) | Breast cysts (PMID: 1387652) |  |
| 10 | 5alpha-pregnan-3beta-ol,20-one sulfate<br>(No dedicated entry, PubChemID 3080643) | Lipid, Estrogenic Steroids | Placenta (PMID: 9449217, 23849932) | Progesterone metabolite, isomer of allopregnanolone (PMID: 16581232) | No currently published evidence directly associating metabolite with specific human genes. |
| 11 | 21-hydroxypregnenolone monosulfate (1)<br>(No dedicated entry, PubChemID 174681) | Lipid, Pregnenolone Steroids | Maternal Plasma (PMID: 31052173) | Spontaneous labor (PMID: 31052173) | No currently published evidence directly associating metabolite with specific human genes. |
| 12 | Decadienedioic acid (HMDB0242172) | Lipid, Fatty Acid, Dicarboxylate | None reported | None reported | No currently published evidence directly associating metabolite with specific human genes. |
| 13 | Pregnenolone/allopregnanolone sulfate (HMDB0240591, HMDB0062782) | Lipid, Progestin Steroids | Placenta (HMDB0240591)<br>Blood, Urine, Hepatic tissue (HMDB0062782) | Spontaneous preterm birth (PMID: 32033212)<br>Metabolic and cardiovascular disorders (PMID: 16902246)<br>Adipose tissue function (PMID: 17374880) | 1. **SULT** – sulfonation (PMID: 37961499)<br>2. **STS** – steroid hormone production (PMID: 24084011)<br>3. **AKR1C4** – biosynthesis of allopregnanolone from progesterone (PMID: 33988248) |
| 14 | Carboxyibuprofen (HMDB0060564) | Xenobiotics, Drug – Analgesics, Anesthetics | Blood<br>Kidney<br>Liver<br>Urine<br>Feces | Pain | **Cytochrome P450 (CYP)** [4] – oxidative metabolism of drugs (PMID: 25502615, 25502615, 37574146) |
| 15 | Dehydrolithocholate (HMDB00502) | Lipid, Secondary Bile Acid Metabolism | Blood<br>Bile<br>Feces<br>Urine<br>Liver<br>Kidney<br>Gallbladder<br>Intestine<br>Gallbladder | Major depressive disorder<br>Alzheimer's<br>Cirrhosis<br>Chronic hepatitis<br>Colorectal cancer | 1. **SLC** [6] – Bile acid and drug transport<br>2. **ABC transporters** [2] – Efflux of bile acids and drugs<br>3. **FABP** [1] – intracellular bile acid transport |
|  |  |  |  | • Chronic Diarrhea<br>• Jaundice |  |
| 16 | Lithocholate<br>(HMDB0000761) | Lipid, Secondary<br>Bile Acid<br>Metabolism | • Blood<br>• Bile<br>• Feces<br>• Urine<br>• Liver<br>• Intestine | • Cystic fibrosis<br>• Cirrhosis<br>• Colorectal cancer | 1. **SLC** [7] – Bile acid and organic anion transport<br>2. **ABC transporters** [2] – Efflux of bile acids and drugs<br>3. **AKR1C** [2] – steroid hormone and bile acid metabolism<br>4. **FABP** [1] – intracellular bile acid transport<br>5. **CYP** [1] – drug and bile acid oxidation<br>6. **NR1H** [1] – regulation of bile acid and cholesterol metabolism |
| 17 | 1-deoxysphinganine<br>(m18:0)<br>(PubMed list) | Lipid,<br>Sphingolipid<br>Synthesis | None reported | None reported | No currently published evidence directly associating metabolite with specific human genes. |
| 18 | N-acetylglucosamine/N-acetylgalactosamine<br>(HMDB0000212, HMDB0000215) | Carbohydrate,<br>Aminosugar<br>Metabolism | • Adipose tissue<br>• Epidermis<br>• Skeletal muscle<br>• Brain<br>• Intestine<br>• Liver<br>• Pancreas<br>• Placenta<br>• Spleen<br>• Testicle<br>• Prostate<br>• Saliva<br>• Feces<br>• Breastmilk<br>• Blood | • Tay-Sachs disease | <u>N-acetylglucosamine</u><br>1. **GALNT/GALT/GALNS** [6] – glycosylation and galactose metabolism<br>2. **B3GNT** [1] – glycan biosynthesis<br>3. **SLC** [1] – nucleotide sugar transport<br>4. **HYAL** [2] – hyaluronic acid degradation<br>5. **SPAM** [1] – cell adhesion and immune signaling<br><u>N-acetylgalactosamine</u><br>1. **LYZ** [5] – antimicrobial activity and cell wall degradation<br>2. **B4GALT** [4] – glycoprotein and glycolipid synthesis<br>3. **HEX** [2] – degradation of glycoproteins and glycolipids<br>4. **CHIA** [2] – degradation of chitin and chitin-like substrates<br>5. **NAG** [2] – cleavage of N-acetylglucosamine residues<br>6. Epimerase/Kinase enzymes (**GNE, RENBP**) [3] – amino sugar metabolism |
|  |  |  |  |  | 7. **NAGK** [1] – phosphorylation of N-acetylglucosamine<br>8. **POMGNT1** [1] – O-linked glycosylation |
| 19 | 5-acetylamino-6-amino-3-methyluracil (HMDB0004400) | Xenobiotics, Xanthine Metabolism | • Blood<br>• Urine<br>• Feces | • Colorectal cancers (PMID: 27275383)<br>• Caffeine metabolism (PMID: 12534641, 15685651) | 1. **NAT** – caffeine metabolism (PMID: 12534641) |
| 20 | Erythrose (HMDB0002649) | Xenobiotics, Food component/plant | • Blood<br>• Cartilage<br>• Feces | • Schizophrenia (PMID: 22007635) | No currently published evidence directly associating metabolite with specific human genes. |
| 21 | 5alpha-pregnan-3beta,20alpha-diol monosulfate (2)<br>(No dedicated entry, 1 measurement identified in GWAS catalog) | Lipid, Progestin Steroids | None reported | None reported | No currently published evidence directly associating metabolite with specific human genes. |
| 22 | N-acetylthreonine (HMDB0062557) | Amino Acid, Glycine, Serine and Threonine Metabolism | • Placenta<br>• Feces | • Colorectal cancers (PMID: 27275383)<br>• Spontaneous preterm birth (PMID: 32033212) | 1. **ACY1** – producing aminoacylase 1 enzyme to break down proteins and release amino acids (PMID: 16465618) |
| 23 | 17beta-estradiol 3-sulfate (HMDB0004448) | Lipid, Estrogenic Steroids | • Blood<br>• Urine<br>• Liver<br>• Kidney | None reported | No currently published evidence directly associating metabolite with specific human genes. |
| 24 | N2,N6-diacetyllysine<br>(No detected entry, PubChemID 7019608) | Amino Acid, Lysine Metabolism | None reported | None reported | No currently published evidence directly associating metabolite with specific human genes. |
| 25 | Ibuprofen (HMDB0001925) | Xenobiotic, Drug – Analgesics, Anesthetics | • Adipose tissue<br>• Epidermis<br>• Skeletal muscle<br>• Blood | • Non-steroidal anti-inflammatory drug<br>• Non-narcotic analgesic<br>• Antipyretic | 1. **AKR1C** [2] – steroid and prostaglandin metabolism<br>2. **UGT** [1] – glucuronidation<br>3. **PTGS** [2] – prostaglandin synthesis (inflammation)<br>4. **CYP2C** [1] – drug metabolism |
|  |  |  | • Cerebrospinal fluid<br>• Bladder<br>• Intestine<br>• Kidney<br>• Pancreas<br>• Placenta<br>• Prostate<br>• Spleen<br>• Testicle<br>• Urine | • Cyclooxygenase 1 & 2 inhibitor | 5. **MAO** [1] – neurotransmitter metabolism<br>6. **ALB** [1] – plasma protein |
| 26 | 4-hydroxybenzoate (HMDB0000500) | Xenobiotic, Benzoate Metabolism | • Blood<br>• Cerebrospinal fluid<br>• Saliva<br>• Urine<br>• Feces<br>• Sweat | • Supradiaphragmatic malignancy | 1. **UGT** [1] – glucuronidation<br>2. **SULT** [1] – sulfonation<br>3. **GLYAT** [1] – acylation (detoxification)<br>4. **COQ** [1] – coenzyme Q biosynthesis |
| 27 | 2-hydroxyibuprofen (HMDB0060920) | Xenobiotic, Drug – Analgesics, Anesthetics | • Blood<br>• Kidney<br>• Liver<br>• Urine<br>• Feces | None reported | 1. **UGT** [1] – glucuronidation |
| 28 | Androstenediol (3beta,17beta) disulfate (HMDB0240313) | Lipid, Androgenic Steroids | • Blood<br>• Placenta<br>• Feces | • Spontaneous preterm birth (PMID: 32033212) | No currently published evidence directly associating metabolite with specific human genes. |
| 29 | 20-hydroxyecdysone (HMDB0030180) | Xenobiotics, Food component/plant | • Feces | • Colorectal cancer (PMID: 25037050) | No currently published evidence directly associating metabolite with specific human genes. |
| 30 | Chiro-inositol (HMDB0240209) | Lipid, Inositol Metabolism | • Urine | None reported | No currently published evidence directly associating metabolite with specific human genes. |

A total of 24 fecal-derived metabolites across seven pathways exhibited significant changes between 24-and 35 weeks gestation **(Table 3)**. Notably, 58.3% increased from 24-and 35 weeks gestation, with sulfate [Xenobiotic, Chemical] (FC = 1.64, p= 0.02), inosine [Nucleotide, Purine Metabolism] (FC = 1.42, p= 0.02), and glycerol 3-phosphate [Lipid, Glycerolipid Metabolism] (FC = 1.41, p= 0.01) demonstrating the largest increases. A total of 41.7% of metabolites decreased during pregnancy, with glycyrrhetinate [Xenobiotics, Food component/plan] (FC = 0.54, p= 0.03), 1-methyl-5-imidazolelactate (FC = 0.55, p= 0.04), and 1-deoxysphinganine (m18:0) [Lipid, Sphingolipid Synthesis] (FC = 0.60, p < 0.001) exhibiting the largest decreases.

**Table 3.** Significant changes in metabolites from 24-to 35 weeks gestation (mid-to-late pregnancy)

|  |  |  |  |  | Associations with PPD |
| --- | --- | --- | --- | --- | --- |
| Pathway (Metabolite count) | Subpathway | Metabolite | Raw fold change value | p-value (q-value) | r (p-value) |
| Lipid (8) | Fatty Acid, Dicarboxylate | 3-methyladipate | 1.22 | 0.0431 (0.8576) | -0.4583 (0.0212)* |
|  | Glycerolipid Metabolism | glycerol 3-phosphate | 1.41 | 0.0141 (0.8576) | -0.0391 (0.8527) |
|  | Sphingolipid Synthesis | 1-deoxysphinganine (m18:0) | 0.55 | 0.0005 (0.5025) | 0.0679 (0.747) |
|  | Sterol | lanosterol | 0.79 | 0.0143 (0.8576) | 0.0158 (0.9403) |
|  | Secondary Bile Acid Metabolism | 3-dehydrodeoxycholate | 0.80 | 0.0475 (0.8576) | 0.2393 (0.2493) |
|  |  | lithocholate | 0.73 | 0.0252 (0.8576) | 0.051 (0.8085) |
|  |  | 12-ketolithocholate | 0.75 | 0.0390 (0.8576) | 0.2654 (0.1998) |
|  |  | dehydrolithocholate | 0.64 | 0.0441 (0.8576) | 0.1977 (0.3434) |
| Nucleotide (6) | Purine Metabolism, (Hypo)Xanthine/Inosine containing | inosine | 1.42 | 0.0224 (0.8576) | -0.1628 (0.4368) |
|  |  | 2'-deoxyinosine | 1.26 | 0.0095 (0.8576) | -0.176 (0.4001) |
|  | Purine Metabolism, Guanine containing | 2'-deoxyguanosine | 1.36 | 0.0460 (0.8576) | -0.0348 (0.869) |
|  | Pyrimidine Metabolism, Uracil containing | 2'-deoxyuridine | 1.24 | 0.0489 (0.8576) | -0.2086 (0.317) |
|  | Pyrimidine Metabolism, Thymine containing | thymidine | 1.23 | 0.0223 (0.8576) | -0.1852 (0.3756) |
|  |  | thymine | 1.17 | 0.0382 (0.8576) | -0.3968 (0.0495)* |
| Xenobiotic (5) | Food component/plant | enterolactone | 0.60 | 0.0116 (0.8576) | -0.2857 (0.1662) |
|  |  | glycyrrhetinate | 0.54 | 0.0260 (0.8576) | 0.0855 (0.6844) |
|  | Bacterial/Fungal | diaminopimelate | 1.23 | 0.0269 (0.8576) | -0.254 (0.2204) |
|  | Drug - Other | N-carbamoylglutamate | 1.36 | 0.0383 (0.8576) | 0.0639 (0.7615) |
|  | Chemical | sulfate | 1.64 | 0.0175 (0.8576) | 0.2285 (0.272) |
| Amino acid (2) | Histidine metabolism | 1-methyl-5-imidazolelactate | 0.55 | 0.0442 (0.8567) | -0.2717 (0.1889) |
|  | Lysine metabolism | N6-formyllysine | 1.31 | 0.0185 (0.8576) | 0.2155 (0.301) |
| Peptide (1) | Dipeptide | alpha-glutamylglutamate | 1.37 | 0.0406 (0.8576) | -0.0681 (0.7465) |
| Carbohydrate (1) | Aminosugar Metabolism | N-acetylmuramate | 1.35 | 0.0033 (0.8576) | -0.4557 (0.0221)* |
| Energy (1) | TCA Cycle | succinylcarnitine (C4-DC) | 0.58 | 0.0100 (0.8576) | -0.1274 (0.5438) |

\*p < 0.05; Fold change > 1 is an increase, < 1 is a decrease; PPD = Postpartum depression as measured by Edinburgh Postnatal Depression Scale scores.

#### Mid-Pregnancy to Postpartum (24 weeks gestation, 6 weeks postpartum)

A total of 26 fecal-derived metabolites across seven pathways exhibited significant changes between 24 weeks gestation and 6 weeks postpartum **(Table 4)**. Three metabolites—5alpha-pregnan-3beta,20alpha-diol monosulfate [Lipid, Progestin steroid] (FC = 3.03, q < 0.001), erythrose [Xenobiotic, Food component/plant] (FC = 0.57, q = 0.01), N-acetylglucosamine/N-acetylgalactosamine [Carbohydrate, Aminosugar metabolism] (FC = 0.56, q = 0.049)—remained significant after FDR. Of the 26 metabolites, 12 (46.2%) increased from 24 weeks gestation to 6 weeks postpartum, while 14 (53.8%) decreased. The most significant increases observed were for N-acetylneuraminate [Amino acid, Polyamine metabolism] (FC = 3.03, p= 0.01), 5-acetylamino-6-amino-3-methyluracil [Xenobiotics, Xanthine metabolism] (FC = 2.32, p= 0.04), and alpha-ketoglutaramate [Amino acid, Glutamate metabolism] (FC = 1.59, p= 0.01), while notable decreases included adipate (C6-DC) [Lipid, Fatty Acid, Dicarboxylate] (FC = 0.30, p= 0.02), 25-hydroxycholesterol sulfate [Lipid, Sterol] (FC = 0.49, p < 0.001), and 3beta-hydroxy-5-cholenoate [Lipid, Secondary bile acid metabolism] (FC = 0.52, p = 0.01).

**Table 4.** Significant changes in metabolites from 24 weeks gestation (mid-pregnancy) to 6 weeks postpartum.

| Pathway<br>(Metabolite<br>count) | Subpathway | Metabolite | Raw fold<br>change<br>value | p-value<br>(q-value) | Pregnancy<br>changes<br>associated<br>with PPD?<br>r (p-value) |
| --- | --- | --- | --- | --- | --- |
| Lipid<br>(9) | Fatty Acid,<br>Dicarboxylate | Adipate (C6-DC) | 1.54 | 0.211<br>(0.5264) | - |
|  |  | heptenedioate (C7:1-DC) | 1.59 | 0.234<br>(0.5264) | - |
|  |  | tetradecanedioate (C14-DC) | 0.57 | 0.0413<br>(0.6424) | - |
|  |  | 2-hydroxysebacate | 1.48 | 0.0342<br>(0.5995) | - |
|  | Endocannabinoid | lignoceroyl ethanolamide<br>(24:0) | 0.57 | 0.0091<br>(0.3370) | - |
|  | Lysophospholipid | 1-stearoyl-GPE (18:0) | 1.18 | 0.0136<br>(0.4406) | - |
|  | Sterol | 25-hydroxycholesterol sulfate | 1.54 | 0.0008<br>(0.0525) | - |
|  | Progestin steroid | 5alpha-pregnan-3beta,20alpha-diol<br>monosulfate (1) | 3.03 | < 0.001<br>(<0.001)* | - |
|  | Secondary bile acid<br>metabolism | 3beta-hydroxy-5-cholenoate | 0.53 | 0.0120<br>(0.4056) | 0.4922<br>(0.0124) |
| Amino acid<br>(8) | Glycine, Serine and<br>Threonine Metabolism | dimethylglycine | 0.86 | 0.0008<br>(0.525) | - |
|  | Glutamate Metabolism | alpha-ketoglutaramate | 0.68 | 0.0093<br>(0.3370) | - |
|  |  | gamma-aminobutyrate<br>(GABA) | 0.58 | 0.0139<br>(0.4406) | - |
|  | Lysine Metabolism | pipecolate | 0.30 | 0.0157<br>(0.4563) | - |
|  | Phenylalanine<br>Metabolism | N-butyryl-phenylalanine | 1.31 | 0.335<br>(0.5995) | - |
|  | Tryptophan<br>Metabolism | 5-hydroxypicolinic acid | 0.72 | 0.498<br>(0.6930) | - |
|  | Leucine, Isoleucine<br>and Valine Metabolism | methylsuccinate | 1.37 | 0.0248<br>(0.5350) | - |
|  | Polyamine Metabolism | N-acetylputrescine | 1.54 | 0.0148<br>(0.4428) | - |
| Xenobiotics<br>(3) | Xanthine Metabolism | 3,7-dimethylurate | 0.73 | 0.0431<br>(0.6424) | - |
|  |  | 5-acetyl-amino-6-amino-3-methyluracil | 0.49 | 0.0399<br>(0.6424) | - |
|  | Food Component/Plant | erythrose | 0.57 | < 0.001<br>(0.0118)* | -0.3983<br>(0.0486) |
| Carbohydrate<br>(3) | Pentose Metabolism | ribitol | 0.52 | 0.0199<br>(0.5264) | - |
|  | Aminosugar<br>Metabolism | N-acetylneuraminate | 1.35 | 0.0261<br>(0.551) | - |
|  |  | N-acetylglucosamine/N-acetylgalactosamine | 0.56 | < 0.001<br>(0.0499)* | - |
| Peptide<br>(1) | Dipeptide | lysylleucine | 1.40 | 0.0452<br>(0.6642) | - |
| Nucleotide<br>(1) | Purine Metabolism,<br>Adenine containing | N6-methyladenosine | 2.32 | 0.393<br>(0.6424) | - |
| Cofactors<br>and Vitamins<br>(1) | Hemoglobin and<br>Porphyrin Metabolism | bilirubin (Z,Z) | 0.59 | 0.0284<br>(0.5747) | - |

\*remained significant after adjusting for multiple comparisons ( $q < 0.05$ ); Fold change $> 1$ is an increase, $< 1$ is a decrease; PPD = Postpartum depression as measured by Edinburgh Postnatal Depression Scale scores.

#### Late-Pregnancy to Postpartum (35 weeks gestation, 6 weeks postpartum)

A total of 55 fecal-derived metabolites across seven pathways exhibited significant changes from 35 weeks gestation to 6 weeks postpartum **(Table 5)**. Four metabolites within the lipid pathway—3-carboxy-4-methyl-5-pentyl-2-furanpropionate [Fatty acid – dicarboxylate] (FC = 2.12, q = 0.002), 5alpha-pregnan-3beta,20alpha-diol monosulfate [Progestin steroid] (FC = 0.80, q = 0.002), lithocholate [Secondary bile acid metabolism] (FC = 1.68, q = 0.008), dehydrolithocholate [Secondary bile acid metabolism] (FC = 1.54, q = 0.03)—remained significant after FDR. Of the 55 metabolites identified, 26 (40%) increased from 35 weeks gestation to 6 weeks postpartum, while 29 (52.7%) decreased. The most significant increases observed were for 5-acetylamino-6-amino-3-methyluracil [Xenobiotic, Xanthine metabolism] (FC = 3.16, p = 0.007), 1-methyl-5-imidazolelactate [Amino acid, Histidine metabolism] (FC = 2.32, p= 0.02), and 3-carboxy-4-methyl-5-pentyl-2-furanpropionate (3-CMPFP) [Lipid, Fatty Acid, Dicarboxylate] (FC = 2.12, p < 0.001). The three metabolites that demonstrated the highest decreases were adipate (C6-DC) [Lipid, Fatty Acid - Dicarboxylate] (FC = 0.29, p= 0.018), gamma-aminobutyrate (GABA) [Amino acid, Glutamate metabolism] (FC = 0.58, p = 0.014), and nicotinate ribonucleoside [Cofactors and vitamins, Nicotinate and Nicotinamide metabolism] (FC = 0.43, p = 0.02).

**Table 5.**
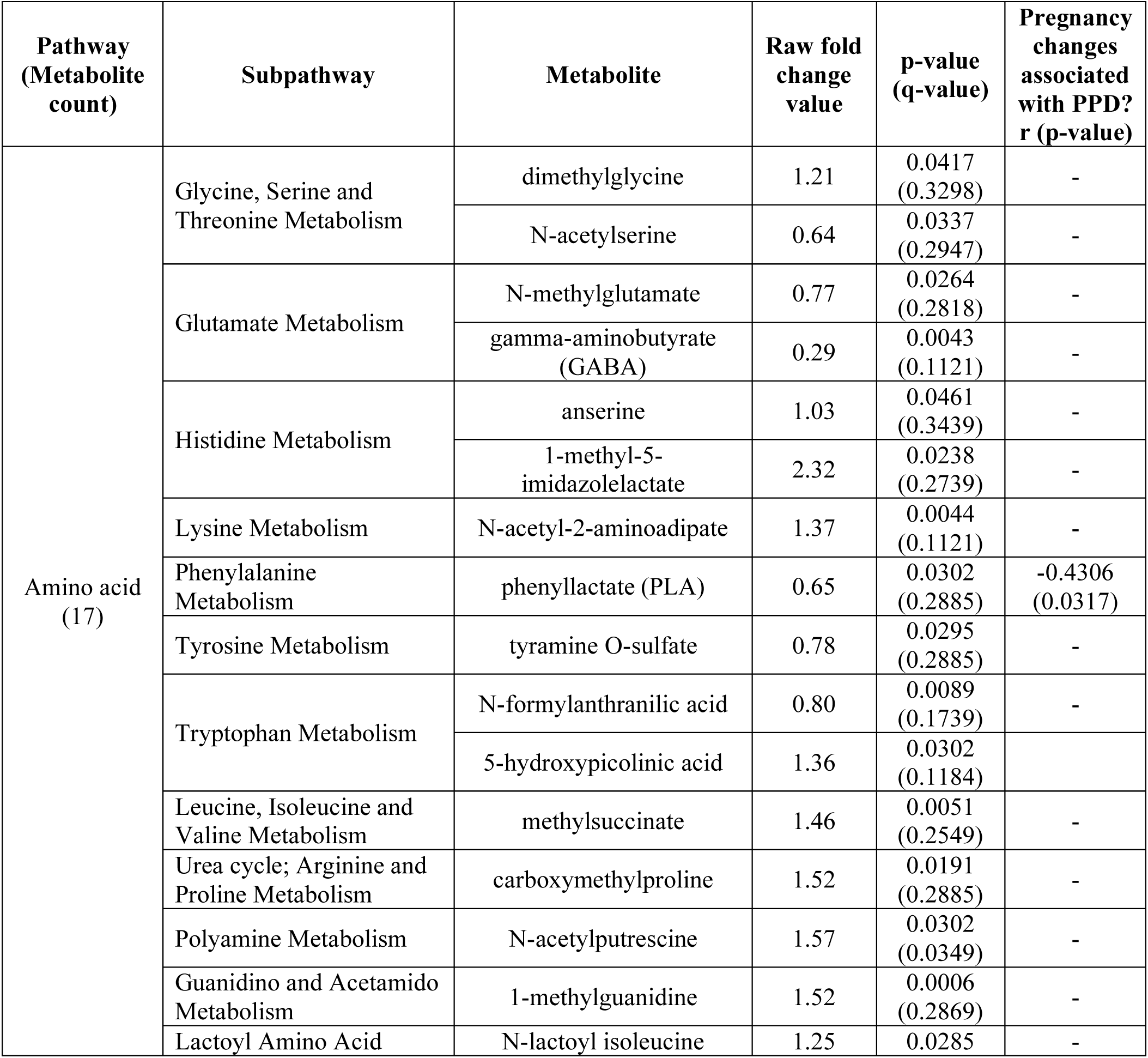
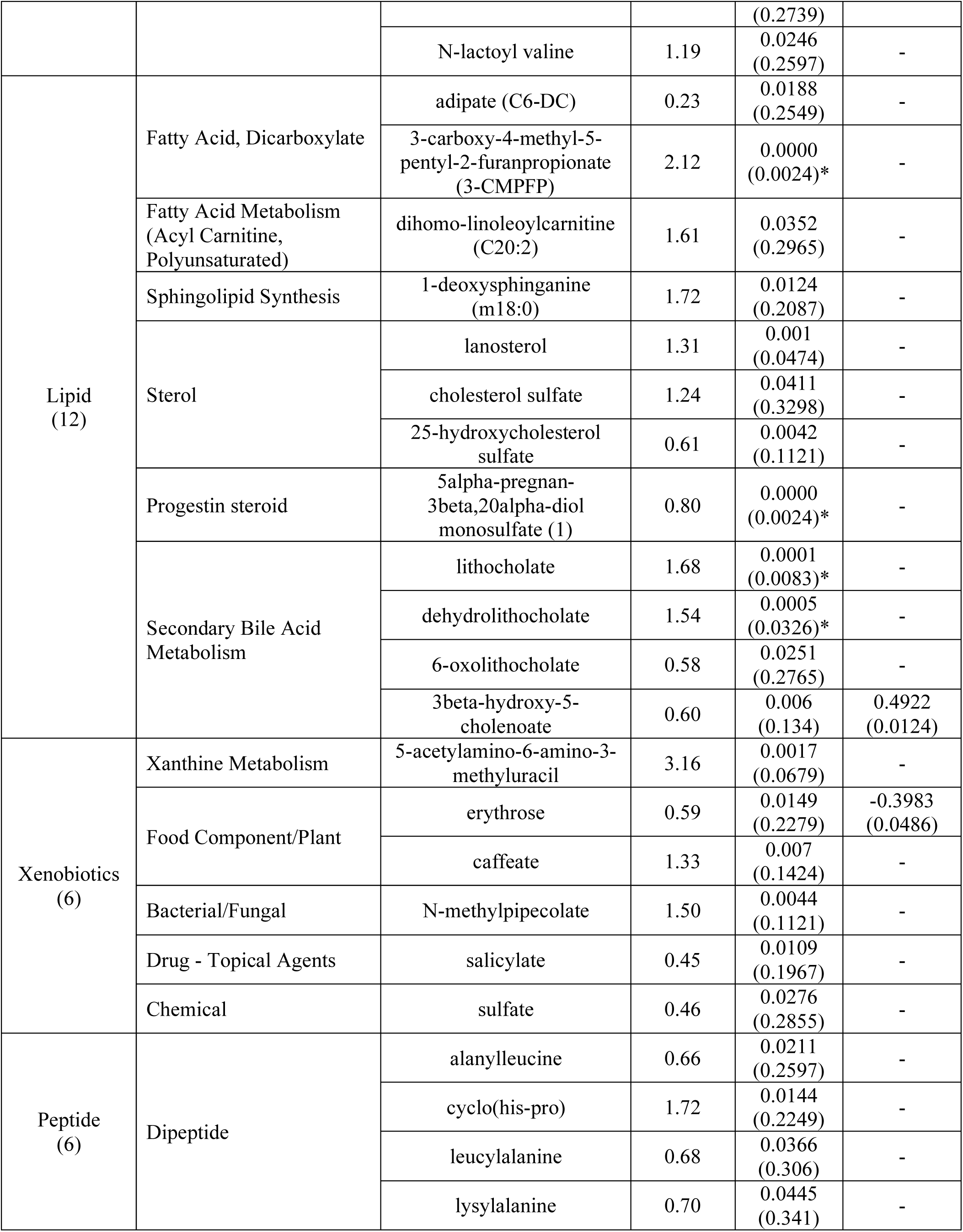
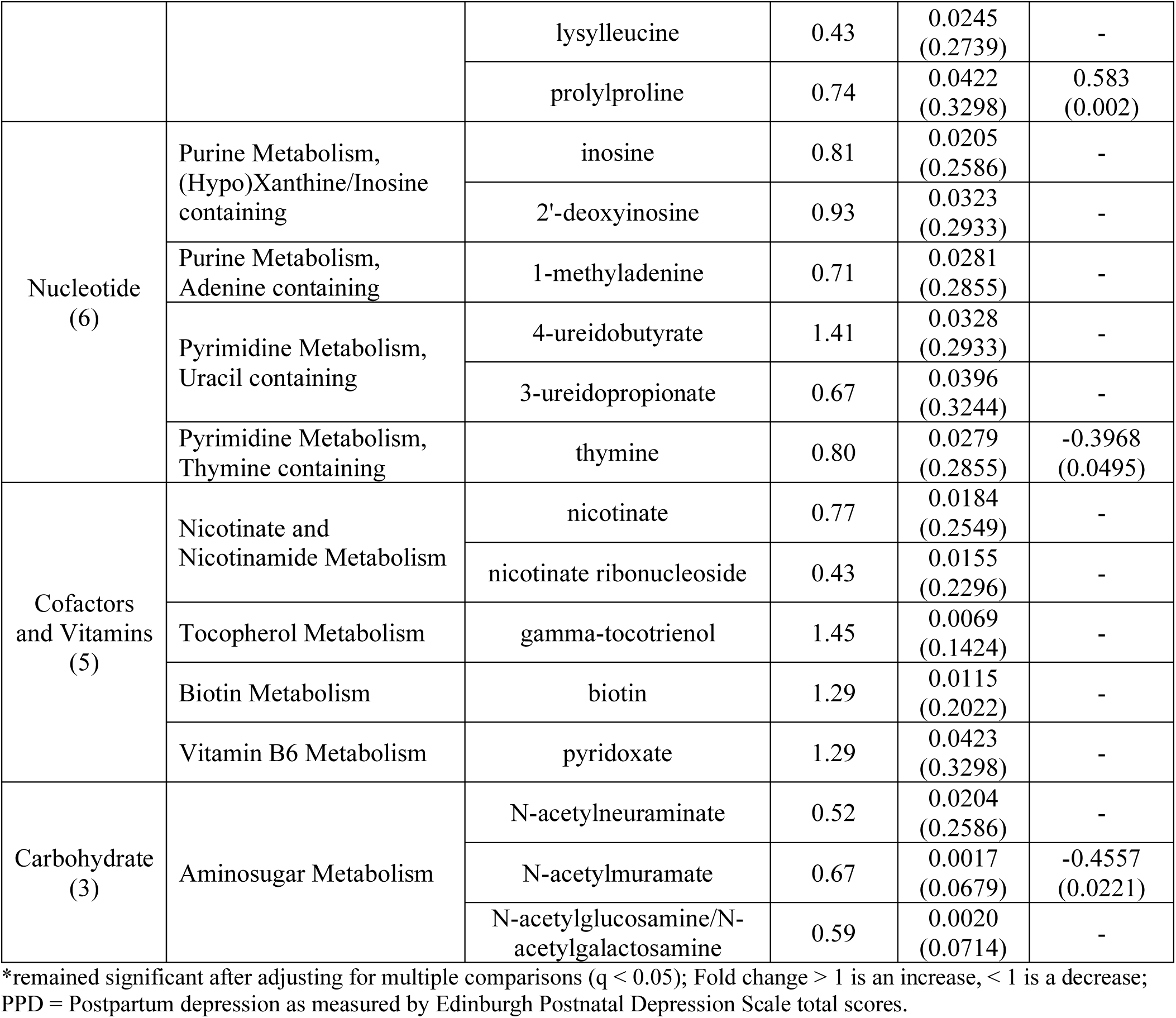
Significant changes in metabolites from 35 weeks gestation (late pregnancy) to 6 weeks postpartum.

### Associations Between Pregnancy Changes in Metabolites and PPD Symptom Severity

A total of 29 pregnancy metabolites across seven pathways were associated with PPD symptom severity at 6 weeks postpartum, with lipid and amino acid metabolites accounting for 58.6% (n = 17) of the associated metabolites **(Table 6)**. A majority of the metabolite changes from mid-to-late pregnancy were moderately associated with PPD symptom severity (82.8%), though 10.3% were trending near moderate and 6.9% near strong. Over half (68.9%) of the metabolite changes from mid-to-late pregnancy that were associated with PPD symptom severity were intermediates compared to microbial byproducts.

**Table 6.** Associations between metabolite changes from mid-to-late pregnancy and postpartum depression symptom severity.

| Pathway<br>(Metabolite<br>count) | Subpathway | Metabolite | r (p-value) | Fold<br>change |
| --- | --- | --- | --- | --- |
| Lipid<br>(10) | Fatty Acid, Dicarboxylate | 3-methyladipate | -0.4583<br>(0.0212) | 1.22* |
|  |  | azelate (C9-DC) | -0.412<br>(0.0407) | 1.02 |
|  | Fatty Acid, Monohydroxy | 3-hydroxyoleate | -0.4<br>(0.0476) | 1.12 |
|  | Fatty Acid, Dihydroxy | 12,13-DiHOME | 0.4128<br>(0.0403) | 0.87 |
|  | Monoacylglycerol | 1-oleoylglycerol (18:1) | 0.5082<br>(0.0095) | 0.31 |
|  |  | 1-linoleoylglycerol (18:2) | 0.4238<br>(0.0347) | 0.32 |
|  | Diacylglycerol | linoleoyl-linoleoyl-glycerol<br>(18:2/18:2) [1] | 0.4792<br>(0.0154) | 1.08 |
|  |  | linoleoyl-linoleoyl-glycerol<br>(18:2/18:2) [2] | 0.5043<br>(0.0101) | 1.15 |
|  | Secondary Bile Acid Metabolism | 3beta-hydroxy-5-cholenoate | 0.4922<br>(0.0124) | 0.86 |
|  |  | isohyodeoxycholate | 0.4824<br>(0.0146) | 1.06 |
| Amino acid<br>(7) | Glutamate Metabolism | N-acetylglutamate | -0.4404<br>(0.0276) | 1.13 |
|  | Histine Metabolism | 4-imidazoleacetate | 0.4218<br>(0.0357) | 0.97 |
|  | Phenylalanine Metabolism | phenyllactate (PLA) | -0.4306<br>(0.0317) | 0.88 |
|  |  | valerylphenylalanine | -0.4041<br>(0.0451) | 0.76 |
|  | Methionine, Cysteine, SAM and<br>Taurine Metabolism | 3-sulfo-alanine | -0.4366<br>(0.0291) | 1.11 |
|  | Urea cycle; Arginine and Proline<br>Metabolism | N-alpha-acetylornithine | -0.3991<br>(0.0481) | 1.00 |
|  | Lactoyl Amino Acid | N-lactoyl phenylalanine | 0.4393<br>(0.028) | 0.51 |
| Xenobiotic<br>(3) | Food Component/Plant | erythrose | -0.3983<br>(0.0486) | 0.99 |
|  |  | 3-indoleglyoxylic acid | 0.4228<br>(0.0353) | 1.08 |
|  |  | 2-aminophenol | -0.5099<br>(0.0092) | 1.14 |
| Carbohydrate<br>(3) | Glycolysis, Gluconeogenesis,<br>and Pyruvate Metabolism | glucose | 0.523<br>(0.0073) | 0.79 |
|  | Pentose Metabolism | ribonate | 0.4942<br>(0.012) | 0.70 |
|  | Aminosugar Metabolism | N-acetylmuramate | -0.4557<br>(0.0221) | 1.35* |
| Peptide<br>(3) | Dipeptide | prolylglycine | 0.5956<br>(0.0017) | 1.05 |
|  |  | prolylproline | 0.583<br>(0.0022) | 1.16 |
|  | Modified Peptides | pyroglutamylalanine | 0.4934<br>(0.0122) | 0.78 |
| Nucleotide<br>(2) | Purine Metabolism, Guanine<br>containing | 7-methylguanine | 0.3996<br>(0.0478) | 0.84 |
|  | Pyrimidine Metabolism,<br>Thymine containing | thymine | -0.3968<br>(0.0495) | 1.17* |
| Cofactors<br>and Vitamins<br>(1) | Ascorbate and Aldarate<br>Metabolism | oxalate (ethanedioate) | 0.4097<br>(0.042) | 1.27 |
| !Other<br>Xenobiotic<br>(1) | Drug - Psychoactive | sertraline | 0.4685<br>(0.0182) | 0.88 |

Only significant correlations ( $p < 0.05$ ) between changes in pregnancy metabolites (24- weeks to 35 weeks gestation) and postpartum depression were reported. Fold change $> 1$ is an increase, $< 1$ is a decrease; Minimal change is $< 0.10$ change; ! indicates fill value $< 80\%$ .

Nearly half (41.4%) of the metabolites significantly associated with PPD symptom severity decreased from mid-to-late pregnancy, while 27.6% exhibited minimal to no change. Only three of the PPD-associated metabolites also showed statically significant changes from mid-to-late pregnancy: 3-methyladipate [Lipid, Fatty acid-Dicarboxylate] (FC = 1.22, p =; r =-0.4583, p = 0.0212), N-acetylmuramate [Carbohydrate, Aminosugar metabolism] (FC = 1.35, p =; r =-0.4557, p = 0.0221), thymine [Nucleotide, Purine metabolism – Thymine containing] (FC = 1.17, p =; r =-0.3968, p = 0.0495). All three metabolites increased from mid-to-late pregnancy and were negatively associated with PPD symptom severity, indicating that greater increases from mid-to-late pregnancy may reflect adaptive metabolic shifts that buffer against PPD symptoms. The remaining 26 metabolites that were significantly associated with PPD symptom severity did not exhibit statistically significant changes from mid-to-late pregnancy. However, 10 out of the 12 (83.3%) metabolites that decreased from mid-to-late pregnancy were positively associated with PPD symptom severity, suggesting greater declines from mid-to-late pregnancy were associated with higher PPD symptom severity. Notably, 40% of those metabolites were involved in lipid metabolism, emphasizing the potential role of dysregulated lipid metabolism during pregnancy in the emergence of PPD symptoms. Interestingly, although only four participants in this study reported taking sertraline (SSRI), sertraline levels decreased from mid-to-late pregnancy and were positively and significantly associated with PPD symptom severity (FC = 0.88; r = 0.4685, p = 0.018), suggesting greater declines in sertraline levels from mid-to-late pregnancy were associated with higher PPD symptom severity.

## DISCUSSION

### Shifts in the Maternal Gut Metabolome

In the current study, we applied a UPLC-MS/MS based untargeted metabolomics approach to maternal fecal samples to examine perinatal shifts in the maternal gut metabolome and associations between changes from mid-to-late pregnancy and PPD symptom severity. We observed differences in metabolic profiles between those with and without a self-reported mood disorder history, suggesting there may be shared alterations in metabolic pathways among those with a history of a mood disorder at some point in their lifetime. Given established links between gut microbiome diversity and metabolic flexibility (de Vos et al., 2022; Edwards et al., 2017; Yao et al., 2020), these results raise the possibility that perinatal individuals with a mood disorder history may experience altered metabolic activity during a life-stage characterized by heightened and prolonged metabolic demands (Kepley, 2023). Since a mood disorder history is a well-established risk factor for PPD (Liu et al., 2022; Yang et al., 2022), these findings may reflect the residual influence a mood disorder can have on metabolic activity, providing a potential biological explanation for the higher incidence of PPD in individuals with a mood disorder history.

Future studies in large, diverse perinatal cohorts are needed to replicate these findings and identify specific pathways and biomarkers driving the observed clustering patterns.

Consistent with evidence from studies utilizing systemic measures (Lindsay et al., 2015; Mitro et al., 2021), the perinatal shifts observed from mid-to-late pregnancy to 6 weeks postpartum reinforce maternal metabolic activity undergoes significant changes after childbirth (Kepley, 2023), marking pregnancy and postpartum are interconnected yet biologically distinct periods. Supporting localized metabolic activity during pregnancy is biologically distinct from that of non-pregnant states (Kepley, 2023), fecal-derived metabolites in our study were observed to distinguish between pregnancy and postpartum timepoints with 96% accuracy. Therefore, this study extends the literature by demonstrating that fecal-derived metabolites not only complement systemic measures (Kazma et al., 2020; Kepley, 2023; Wishart, 2019) but also remain relatively stable from mid-to-late pregnancy, despite extrinsic influences—a common criticism raised regarding the reliability of gut-derived measures. Collectively, these findings highlight the critical importance of investigating perinatal health and disease within the physiological contexts of pregnancy and postpartum, and cautions against over-reliance on evidence derived from non-perinatal populations or those without reproductive organs in informing perinatal health care and targeted approaches in perinatal research.

Mid-pregnancy is characterized by anabolic processes that promote nutrient storage and tissue growth, while late pregnancy and postpartum mark a transition to a catabolic state in which stored nutrients are mobilized to support maternal-fetal health and lactation (Kazma et al., 2020; Kepley, 2023). We observed metabolite excretion to be lowest from mid-to-late pregnancy but double from late pregnancy to postpartum, aligning with these physiological transitions. Further consistent with studies examining systemic measures, lipid and amino acid metabolism emerged as core pathways across perinatal timepoints (Lindsay et al., 2015; Wishart, 2019). The concordance between systemic measures (Bränn et al., 2021; Kimmel et al., 2022; Lindsay et al., 2015; Mitro et al., 2021) and the fecal-derived metabolites observed in this study suggest coordinated metabolic adaptations occur across biological systems, highlighting the need for future research applying multi-omics methods to integratively examine localized (gut) and systemic biomarkers to advance understanding of the biological networks underlying perinatal health and disease (e.g., PPD).

### Associations with PPD

A majority (70%) of pregnancy metabolite changes that were associated with PPD symptom severity were intermediates. Unlike metabolic byproducts, which are typically excreted as waste, intermediates are transient, bioactive compounds formed within active metabolic pathways and play a regulatory role in maintaining metabolic homeostasis (Alonso et al., 2015; Wishart, 2019; Xu et al., 2019). Thus, examining metabolic intermediates may be useful for evaluating how pregnant individuals are adapting to the metabolic demands of pregnancy, potentially revealing early perturbations within biological pathways that contribute to PPD vulnerability. Notably, greater declines in metabolites from mid-to-late pregnancy—nearly 40% of which were lipid metabolites—were associated with higher PPD symptom severity, while increases appeared protective. Lipid-derived metabolites are not merely byproducts of fat metabolism. Many function as signaling molecules, with some such as steroid hormones, playing a critical role in mood regulation by modulating neurotransmitter receptors (Deligiannidis et al., 2021, 2023; Lindsay et al., 2015). Zuranolone, a synthetic neurosteriod (allopregnanolone) analog, alleviates PPD symptoms by enhancing the activity of GABAA receptors (Deligiannidis et al., 2021, 2023). While further research is needed, our findings support the mechanistic rationale underlying Zuranolone’s therapeutic action with respect to lipid metabolism involvement, but extends the literature by identifying pregnancy-specific, localized changes in metabolic activity that may underly pathophysiological trajectories leading to PPD. Additionally, though pregnancy changes in fecal-derived GABA were not associated with PPD, it was one of three metabolites demonstrating significant decreases from late-pregnancy to postpartum. These findings warrant further investigation, as this study did not assess whether metabolite changes from late pregnancy to postpartum were associated with PPD symptom severity. Future studies should consider associations between fecal-derived metabolites at other perinatal timepoints and PPD symptom severity to better characterize how shifts in GABA and other key lipid and amino acid metabolites relate to PPD symptom severity.

Several of the fecal-derived metabolites associated with PPD—such as 3-methyladipate and adipate (lipid metabolism), GABA and alpha-ketoglutaramate (amino acid metabolism), and glycyrrhetinate, erythrose (xenobiotic metabolism)—though most are endogenously derived, can be modified by extrinsic (e.g., diet, medication) and intrinsic (e.g., gut microbiota) factors (Braga et al., 2024; Ohue-Kitano et al., 2024; Strandwitz et al., 2019; TeSlaa et al., 2023). Downstream-omics biomarkers (e.g., microbiome, metabolome) are recognized for their responsiveness to environmental, social, and behavioral influences (Graw et al., 2021; Hasin et al., 2017; Sun & Hu, 2016). While this sensitivity has raised concerns within the broader scientific community about their reliability for biomarker discovery and clinical translation (Estrela et al., 2025), it also highlights their unique strength in measuring dynamic, context-dependent biological processes. This makes them particularly well-suited in the study of complex conditions like PPD, where risk is multifactorial and shaped by both the unique physiological contexts of this lifestage and variable extrinsic influences (Dagher et al., 2021; Kepley, 2023). Therefore, rather than viewing this context sensitivity as a limitation, this sensitivity can be leveraged to measure biological responses to modifiable exposures (e.g., pregnancy symptoms, dietary intake, social determinants), potentially uncovering more comprehensive and actionable markers of risk (Sun & Hu, 2016). As such, future research should prioritize integrative approaches that combine downstream-omics data with contextual assessments, enabling a multidimensional understanding of PPD that can inform personalized strategies for risk identification and multi-level interventions (e.g., medication, support groups, increased food access, policy changes).

Lastly, while the observed association between sertraline exposure and PPD symptoms may be spurious due to the limited sample size, it raises the prospect of leveraging metabolomics to advance understanding of psychopharmacological responses during a period of profound metabolic adaptations. Additional research is needed to evaluate the potential of fecal metabolomic profiling as a non-invasive tool for monitoring responses to pharmacological interventions, which could support tailored treatment approaches to optimize outcomes for both mother and child.

### Limitations

First, as a secondary analysis, this study is limited to the participant characteristics, variables, measures, and timepoints collected as part of the parent study. Second, we utilized samples from an ongoing prospective cohort study that followed mother-child dyads from early pregnancy up to two years postpartum. As a result, the sample size was limited to the 25 participants with complete datasets across all relevant timepoints at the time of analysis. Third, the participant demographics were relatively homogenous, limiting generalizability but offering a valuable benchmark for future comparisons among more diverse sociodemographic groups. Fourth, without a clear understanding of metabolite origin—whether endogenous (host-derived) or exogenous (microbial or dietary)—it remains difficult to determine the extent to which the observed changes reflect intrinsic physiological shifts versus physiological responses to extrinsic influences. Future studies integrating both localized (microbiome, metabolome) and systemic biomarkers (metabolome, other circulating-omics markers) are needed to disentangle these contributions and advance understanding of maternal metabolic adaptations during pregnancy and their role in shaping PPD risk. Lastly, due to the paucity of literature on fecal-derived metabolites in perinatal populations and the underutilization of downstream-omics measures in PPD research, this study is intended to be hypothesis-generating rather than hypothesis-testing.

## CONCLUSION

This study establishes the feasibility and utility of applying untargeted metabolomics (UPLC-MS/MS) to fecal samples to longitudinally investigate microbiome-host interactions across the perinatal continuum, offering localized insights into metabolic activity and associations with PPD symptom severity. Lipid, amino acid, and xenobiotic metabolism emerged as core pathways contributing to perinatal shifts in the maternal gut metabolome, with changes in lipid and amino acid metabolites from mid-to-late pregnancy accounting for over half of the metabolites associated with PPD symptom severity. Findings highlight pregnancy as a potential window for identifying risk biomarkers and reinforces the notion that PPD likely arises from perturbations within a biological network, rather than disruptions in any single system or pathway. Therefore, the ultimate contribution of this study lies in introducing an underutilized-omics layer to locally measure host-microbiome interactions in perinatal populations, laying the groundwork for future multi-omics research integrating localized (gut) and systemic biomarkers to deepen understanding of the biological networks underlying PPD.

## STATEMENTS AND DECLARATIONS

### Conflicts of Interest Statement

There are none to disclose.

### Author’s contributions

KDL was responsible for conceptualization, data analysis, validation, visualization, and writing – original draft; MLW contributed as a supervisor to KDL as well as writing – review and editing; KL contributed to writing – review and editing; SG and OFR contributed to data curation and writing – review and editing; SD contributed to parent study project administration, data collection, writing – review and editing; EF contributed to writing – review and editing; and EMW, PI of the parent study, contributed as a supervisor to all research staff throughout the parent study and KDL in the present study, as well as project administration, resources, and writing – review and editing.

### Data and Software Availability

The de-identified data used and analyzed in this study will only be shared from the principal investigator (PI) of the parent study (EMW) upon reasonable request. There is no software to make available as a result of this study.

### Ethics Approval and Consent to Participate

This study was conducted in accordance with the Declaration of Helsinki and approved by the Institutional Review Board (2018-05-0127) at the University of Texas at Austin. Written informed consent was obtained from all participants in the study. Participation was voluntary, and participants were provided with detailed information about the study’s objectives, procedures, potential risks, and benefits before consenting.

### Funding Information

This project was supported by the National Institutes of Health [NICHD 5R00HD086304-06] (EMW), [T32NR019035] (KDL), and a Population Research Center (PRC) Seed Grant at The University of Texas at Austin [P2CHD042849, NICHD] (MLW). The content is solely the responsibility of the authors and does not necessarily represent the official views of the National Institutes of Health or any affiliated institutions.

## Acknowledgments

We extend our sincere gratitude to all the participants, clinical and research staff, and collaborators whose valuable contributions continue to enable progress in advancing maternal mental health care.

## Clinical Trial Number of Parent Study

NCT04132310; https://www.clinicaltrials.gov/; Registration date: 10/18/2019

## Notes

### Competing Interest Statement

The authors have declared no competing interest.

